# Cell-free DNA as a biomarker for prostate cancer: elevated concentration and decreased fragment size

**DOI:** 10.1101/2020.07.26.215020

**Authors:** Emmalyn Chen, Clinton L. Cario, Lancelote Leong, Karen Lopez, César P. Márquez, Carissa Chu, Patricia S. Li, Erica Oropeza, Imelda Tenggara, Janet Cowan, Jeffry P. Simko, June M. Chan, Terence Friedlander, Alexander W. Wyatt, Rahul Aggarwal, Pamela L. Paris, Peter R. Carroll, Felix Feng, John S. Witte

## Abstract

**Purpose:** Prostate cancer is the most commonly diagnosed neoplasm in American men. Although existing biomarkers may detect localized prostate cancer, additional strategies are necessary for improving detection and identifying aggressive disease that may require further intervention. One promising, minimally invasive biomarker is cell-free DNA (cfDNA), which consist of short DNA fragments released into circulation by dying or lysed cells that may reflect underlying cancer. Here we investigated whether differences in cfDNA concentration and cfDNA fragment size could improve the sensitivity for detecting more advanced and aggressive prostate cancer.

**Materials and Methods:** This study included 268 individuals: 34 healthy controls, 112 men with localized prostate cancer who underwent radical prostatectomy (RP), and 122 men with metastatic castration-resistant prostate cancer (mCRPC). Plasma cfDNA concentration and fragment size were quantified with the Qubit 3.0 and the 2100 Bioanalyzer. The potential relationship between cfDNA concentration or fragment size and localized or mCRPC prostate cancer was evaluated with descriptive statistics, logistic regression, and area under the curve analysis with cross-validation.

**Results:** Plasma cfDNA concentrations were elevated in mCRPC patients in comparison to localized disease (OR_5 ng/mL_ = 1.34, P = 0.027) or to being a control (OR_5 ng/mL_ = 1.69, P = 0.034). Decreased average fragment size was associated with an increased risk of localized disease compared to controls (OR_5bp_ = 0.77, P = 0.0008).

**Conclusion:** This study suggests that cfDNA concentration and average cfDNA fragment size may provide a quick, cost-effective approach to help determine which patients will benefit most from further screening and/or disease monitoring to help improve prostate cancer outcomes.

## Introduction

Prostate cancer accounts for approximately 20% of all new cancer diagnoses in American men. While individuals diagnosed with localized disease have a 98% 5-year survival rate, an estimated 33,330 men will die from aggressive and metastatic disease in 2020^1^. There are a number of existing biomarkers routinely used for prostate cancer diagnosis and monitoring, including prostate-specific antigen (PSA), PHI, 4Kscore, PCA3 expression, parametric MRI, and hypermethylation of GSTP1, APC, and RASSF1^2,3^. These have varying levels of sensitivity and specificity, and additional biomarkers for prostate cancer are necessary to reduce over-diagnosis and over-treatment of this common, but complex disease.

Cell-free DNA (cfDNA) is a promising, minimally invasive biomarker that may originate from cell lysis, apoptosis, necrosis, and active release of DNA fragments into circulation^4–7^. In healthy individuals, cfDNA is predominantly of hematopoietic origin^8^. In cancer patients, cfDNA includes DNA of hematopoietic origin, as well as circulating tumor DNA (ctDNA) derived from tumor cells. Two commonly used methods to profile cfDNA are: 1) quantification of cfDNA based on spectrophotometry, electrophoresis, or quantitative PCR (qPCR); and 2) genomic interrogation of ctDNA fragments with next-generation sequencing, BEAMing (beads, emulsion, amplification, and magnetics), or droplet digital PCR (ddPCR).

While genomic interrogation allows for the detection of cancer-specific fragments, this can have a number of challenges (e.g., sufficient cfDNA, sequencing depth, and mutation panel selection). In contrast, quantification of overall cfDNA concentrations and assessment of cfDNA fragment size may provide a quick, cost-effective method in addition to other biomarkers such as PSA, and deliver insight into whether a patient should undergo further biopsy and potentially genomic testing.

Elevated concentrations of cfDNA were initially reported in patients with leukemia and autoimmune disease^9,10^. Subsequent studies have also determined that high concentrations of cfDNA are typically associated with poor survival in several cancers^11,12^. For prostate cancer, increased plasma cfDNA concentrations were found in patients with lymph node and distant metastases^13^. Elevated preoperative serum cfDNA concentrations in men with localized prostate cancer who underwent RP have been associated with PSA recurrence, independent of surgical margin and lymph node status, as well as Gleason score and pathologic stage^14^.

In addition to overall cfDNA concentrations, cfDNA fragment size may provide diagnostic and prognostic value. DNA integrity, which measures the ratio of all cfDNA fragments (ALU 247bp) to shorter fragments (ALU 115bp) has distinguished prostate cancer from benign prostatic hyperplasia (BPH)^15^. Pre-treatment cfDNA concentration and fragment size were predictive of advanced pancreatic cancer progression-free survival and overall survival^16^. Furthermore, tumor fragments in cfDNA appeared shorter in size than fragments that originated from non-malignant cells^17–19^.

Here, we evaluate whether baseline plasma cfDNA concentrations and cfDNA fragment size can differentiate among: 1) men with prostate cancer and controls; and 2) clinical characteristics or biochemical recurrence among men with localized disease (i.e., PSA at diagnosis, Gleason, organ confinement, extraprostatic extension, seminal vesicle invasion, lymph node invasion, and RNA gene expression). While cfDNA concentration data was available for mCRPC and localized prostate cancer groups, cfDNA fragment size data was only available for patients with localized disease.

## Materials and Methods

### Patient Cohort

From August 2015 to November 2019, biological samples from a total of 268 individuals were included in this study: 34 healthy donors, 112 patients with localized prostate cancer, and 124 mCRPC patients (Table 1). Twenty-eight healthy donor samples were obtained from StemCell (StemCell Technologies, Seattle, WA), and six healthy samples were collected from volunteers at UCSF. All patients with localized disease underwent radical prostatectomy (RP) at UCSF, and 35/112 patients had a Decipher score of RNA gene expression available (GenomeDx Biosciences, Vancouver, British Columbia, Canada)^20^. For the patients with localized disease, blood samples were collected the day of surgery before RP, and for five of these men blood samples were collected two months after surgery. Clinicopathologic variables that play an important role in surgical management after prostatectomy were also collected, including clinical T stage, pathologic Gleason score, preoperative PSA, and risk prediction models including the Cancer of the Prostate Risk Assessment Postsurgical (CAPRA-S) score and the Decipher score^21,22^. Known predictors of biochemical recurrence (BCR), including organ confinement, extraprostatic extension, seminal vesicle invasion, and lymph node involvement were also identified^23^. Biochemical recurrence was defined as two consecutive PSA levels of ≥0.2 ng/mL eight weeks after surgery.

**Table 1.** Clinical characteristics of individuals included in the study at baseline.

|  | Healthy<br>N = 34 | Localized<br>N = 112 | mCRPC<br>N = 122 |
| --- | --- | --- | --- |
| **Age (years)** |  |  |  |
| Median ± IQR | 60 ± 17 | 65 ± 10 | 71 ± 9 |
| Range | 41 – 74 | 43 – 78 | 47 – 91 |
| **Pathologic Gleason\*** |  |  |  |
| 6 | - | 6 | - |
| 7 | - | 75 | - |
| 8-10 | - | 30 | - |
| **Pathologic Stage** |  |  |  |
| Organ confined (pT2) | - | 46 | - |
| Not organ confined (pT3, pT4) | - | 66 | - |
| Extraprostatic extension (pT3a) | - | 43 | - |
| Seminal vesicle invasion (pT3b) | - | 15 | - |
| Lymph node involvement (N1) | - | 17 | - |
| **PSA (ng/mL)** |  |  |  |
| Median ± IQR | - | 6.9 ± 7.4 | 50.8 ± 128 |
| Range | - | 1.21 – 70.0 | 0 – 3007 |
| **cfDNA Concentration (ng/mL)<sup>†</sup>** |  |  |  |
| Median ± IQR | 7.9 ± 4.0 | 6.7 ± 5.8 | 13.8 ± 28.1 |
| Range | 0.29 – 16.9 | 1.22 – 53.9 | 1 – 1380 |
| **cfDNA Fragment Size (bp)** |  |  |  |
| Median ± IQR | 177.5 ± 29.5 | 173 ± 6 | - |
| Range | 142 – 265 | 135 – 280 | - |

\*One man with unknown data in the cohort.
<sup>†</sup> Concentration data collected on a subset of individuals (31 healthy, 45 localized, and 122 mCRPC).

While most of the cohort was collected prospectively, a subset of 110 mCRPC patients were recruited through the Stand Up 2 Cancer/Prostate Cancer Foundation-funded West Coast Prostate Cancer Dream Team Project (IRB 12-10340). Fourteen mCRPC patients were recruited through UCSF^24^. For mCRPC patients, blood samples were collected prior to treatment initiation. Clinicopathologic characteristics were collected for all patients (Table 1). Approval for this study was granted by the local ethics review board (IRB 11-05226 and IRB 12-09659), and written informed consent was obtained from all patients.

### cfDNA Extraction from Blood

For healthy controls, whole peripheral blood samples were collected from individuals in PAXgene Blood ccfDNA tubes (Qiagen, Redwood City, CA). Healthy samples collected by StemCell were shipped at room temperature, arriving within 7 days for sample processing. Whole peripheral blood samples were collected immediately before surgery for patients with localized disease or at the time of follow-up and before treatment initiation for mCRPC patients. Plasma was generated from whole blood samples within 2 hrs for blood collected in K3EDTA tubes or within 7 days for blood collected in PAXgene Blood ccfDNA tubes with a two-step centrifugation protocol: first centrifuging the blood at 1,900g for 10 min at 21°C, followed by centrifugation of the supernatant at 16,000g for 10 min to remove leukocytes and cellular debris. DNA was extracted from 7 to 55 mL of plasma using the Qiagen QIAamp Circulating Nucleic Acid Kit (Qiagen, Redwood City, CA), and double eluted with 40 μL of Qiagen Elution Buffer. Extracted DNA was stored at -20°C prior to further analysis.

### cfDNA Fragment Size and Concentration

Extracted DNA was quantified with a Qubit 3.0 Fluorometer and a DNA dsDNA HS Assay Kit (Life Technologies, Carlsbad, CA), as well as on the 2100 Bioanalyzer with High Sensitivity DNA Chips (Agilent Technologies, Santa Clara, CA) for assessment of sample purity, concentration, and fragment size distribution according to the manufacturer’s instructions. The average fragment size was determined with the Agilent 2100 Bioanalyzer Expert software, and calculated across the first three peaks 75–675bp corresponding to the length of nucleosomal footprints and linkers derived from apoptotic cells (Supplementary Figure 1). The final plasma cfDNA concentrations were calculated by adjusting for the initial plasma and final elution volumes, and quantified with a Qubit 3.0 for a subset of patients (Supplementary Table 1). Assessment of cfDNA fragment size and concentration was performed without prior knowledge of clinical data. Average cfDNA fragment size was not available for mCRPC patients, since samples were not run on the 2100 Bioanalyzer.

### Statistical Analysis

Our primary analysis assessed the relationship between cfDNA concentration or average fragment size and prostate cancer, comparing three groups: healthy controls, men with localized disease, and men with mCRPC. Here we used descriptive statistics, logistic regression, and receiver operating characteristic (ROC) curves. Since cfDNA concentration and average fragment size were not normally distributed (P < 0.001, Shapiro–Wilk test), we evaluated the difference in descriptive statistics across prostate cancer diagnoses using the Mann-Whitney non-parametric test. We also evaluated differences in cfDNA concentration quantified between 90–150bp, which is known to be enriched for circulating tumor DNA fragments specifically^17^. Then, we further investigated the potential relationship between cfDNA concentration and prostate cancer diagnoses using logistic regression models (crude, and then adjusting for age at time of blood draw and baseline PSA when available). The ability of cfDNA concentration to discriminate between prostate cancer diagnoses was further assessed based on the area under the curve (AUC) from a Receiver Operating Characeristics (ROC) curve analysis with k-fold cross-validation (a minimum of ten observations per fold) and bootstrap resampling (n = 100). These analyses were also performed with average cfDNA fragment size to distinguish patients with localized disease from controls.

We also undertook secondary analyses investigating the relationship between baseline cfDNA concentration or fragment size and clinical characteristics among patients with localized disease. For continuous characteristics, comparisons were made using cfDNA concentration and Pearson correlation coefficients (i.e., age at diagnosis, PSA at diagnosis, Decipher score, time to salvage therapy, average cfDNA fragment size, and postoperative CAPRA-S score). For categorical clinical features, we assessed the potential relationship between log-transformed cfDNA concentration (for normality) and other clinical features (pathologic Gleason score, organ confinement, extraprostatic extension, seminal vesicle invasion, pathologic lymph node status, biochemical recurrence, and clinical T stage) with Student’s t–tests. Mann-Whitney U-tests were used to assess the association between the average cfDNA fragment size and the same clinical features. As with localized disease, we also evaluated the relationship between cfDNA concentration and age at time of blood draw for healthy individuals and mCRPC patients. Finally, we evaluated the association between cfDNA concentration or fragment size and biochemical recurrence-free survival with Cox proportional hazards models for patients with localized disease. All data analyses were performed using R version 3.6.1.

## Results

### cfDNA Concentration and Prostate Cancer

The median cfDNA concentration was 7.9 ng/mL (IQR, 4.0 ng/mL) for controls, 6.7 ng/mL (IQR, 5.8 ng/mL) for patients with localized disease, and 13.8 ng/mL (IQR, 28.1 ng/mL) for patients with mCRPC (Table 1; Figure 1). The average cfDNA levels in mCRPC patients were statistically significantly higher than those observed in controls (P < 0.0001) or those with localized prostate cancer (P < 0.0001).

**Figure 1.**
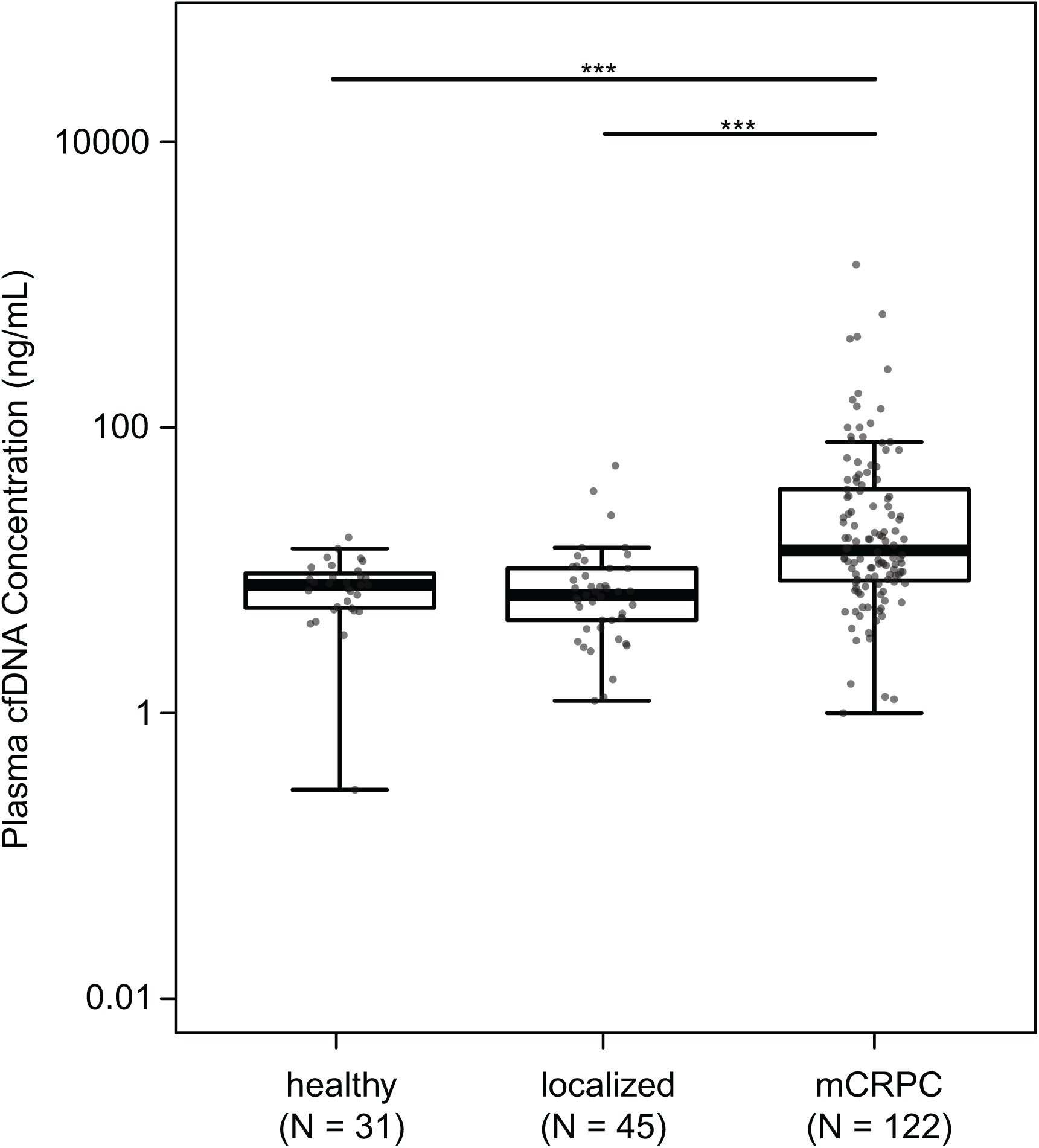
Distribution of plasma cfDNA concentration in healthy individuals, patients with localized disease, and patients with mCRPC. Boxplots and points identify the minimum, interquartile range, median, and maximum values for each group. The Mann-Whitney test was applied to test differences in cfDNA concentration between groups. *** P < 0.0001

These observations were further supported by results from the logistic regression models, including those adjusting for age and PSA levels (Table 2). A 5 ng/mL increase in cfDNA concentration was positively associated with mCRPC in comparison to localized disease (OR_crude_ = 1.47, P = 0.0017; OR_adjusted_ = 1.34, P = 0.027) or to being healthy (OR_crude_ = 1.93, P = 0.0025; OR_adjusted_ = 1.69, P = 0.034). Plasma cfDNA concentration was not associated with having localized disease in comparison to healthy individuals (OR_crude_ = 1.10, P = 0.64; OR_adjusted_ = 1.05, P = 0.72).

**Table 2.** Association between increase in cfDNA concentration (5 ng/mL) or in cfDNA fragment size (5bp) and prostate cancer status. Results from crude univariate and adjusted multivariate logistic regression analyses (adjusted for age, and PSA when available).

| cfDNA Measure | Prostate Cancer Status | Crude |  | Adjusted |  |
| --- | --- | --- | --- | --- | --- |
|  |  | Odds Ratio<br>(95% CI) | P-value | Odds Ratio<br>(95% CI) | P-value |
| cfDNA Concentration<br>(5 ng/mL) | **Localized vs Healthy** | 1.10<br>(0.82 – 1.61) | 0.64 | 1.05<br>(0.77 – 1.69)* | 0.72 |
|  | **mCRPC vs Healthy** | 1.93<br>(1.34 – 3.18) | 0.0025 | 1.69<br>(1.16 – 2.93)* | 0.034 |
|  | **mCRPC vs Localized** | 1.47<br>(1.22 – 2.01) | 0.0017 | 1.34<br>(1.05 – 1.76)<sup>†</sup> | 0.027 |
| cfDNA Fragment Size<br>(5bp) | **Localized vs Healthy** | 0.86<br>(0.73 – 0.9) | 0.003 | 0.77<br>(0.66 – 0.90)* | 0.0008 |

\* Adjusted for age
<sup>†</sup> Adjusted for age and PSA

In our ROC curve analysis, plasma cfDNA concentration was able to distinguish between mCRPC patients from healthy individuals and those with localized disease (Figure 2), with an estimated AUC of 0.83 (95% CI, 0.72–0.91) and 0.81 (95% CI, 0.74–0.87), respectively.

**Figure 2.**
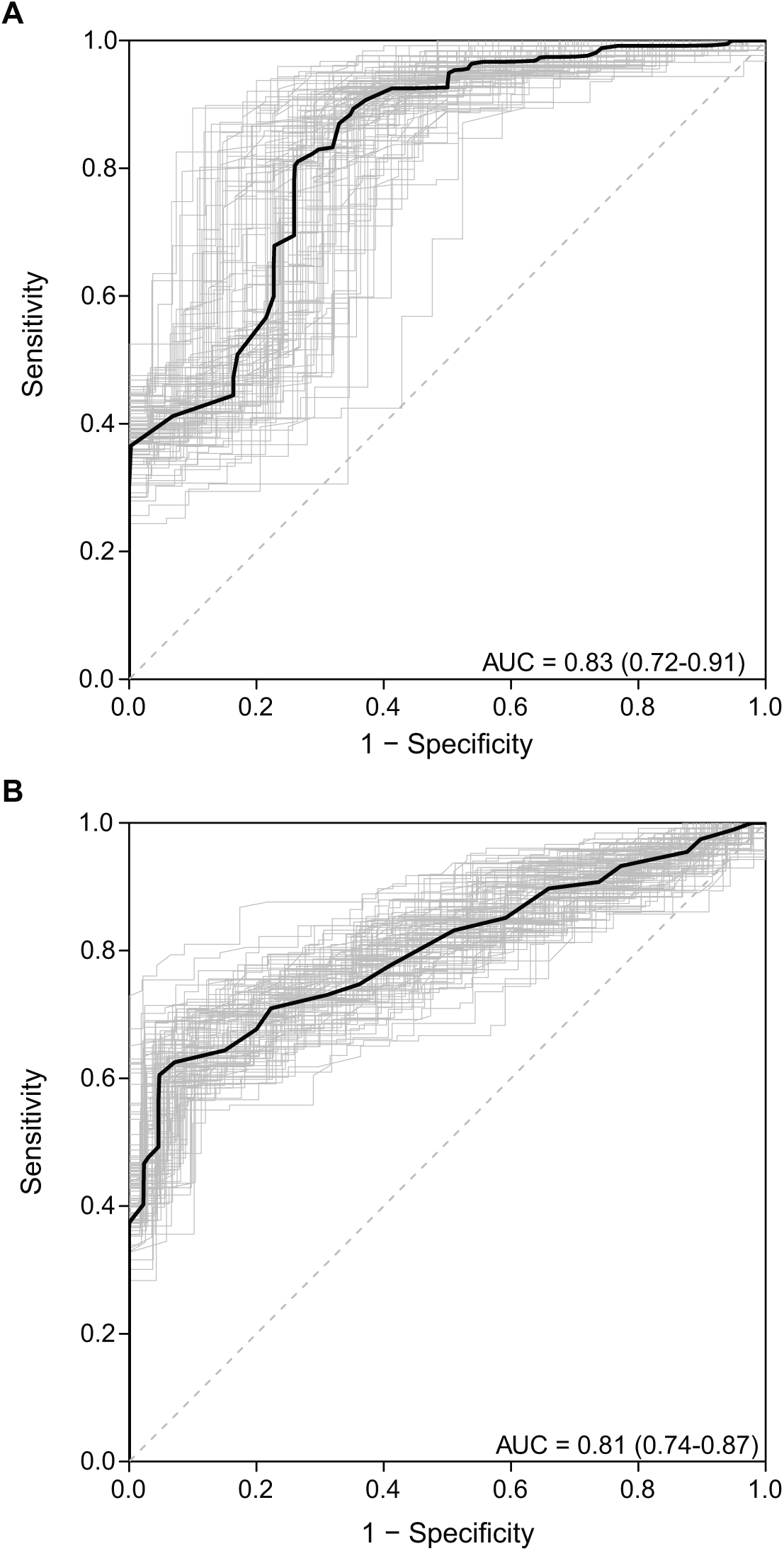
Receiver operating characteristic (ROC) curves for cfDNA concentration comparison between A) healthy individuals and mCRPC, and B) patients with localized disease and mCRPC. Area under the curve (AUC) and 95% CI were estimated with k-fold cross-validation and bootstrap resampling.

### cfDNA Fragment Size and Prostate Cancer

The median of the average cfDNA fragment size in patients with localized disease was 173bp (range, 135-280bp), and in controls was 177.5bp (range, 142-265bp) (Table 1). This lower average cfDNA fragment size in patients with localized disease was statistically significantly different from that observed in controls (P = 0.0009, Figure 3). Results from the logistic regression analysis further indicate that average fragment size was inversely associated with localized prostate cancer (in comparison to healthy individuals): for a 5bp increase in fragment size, the OR_crude_ = 0.86 (P = 0.003) and OR_adjusted_ = 0.77 (P = 0.0008; Table 2). The estimated ROC AUC for distinguishing between healthy individuals and patients with localized prostate cancer using average cfDNA fragment size was 0.64 as defined by k-fold cross-validation. There was no difference in cfDNA concentration quantified across 90–150bp between healthy individuals and patients with localized disease (Supplementary Figure 4)^17^.

**Figure 3.**
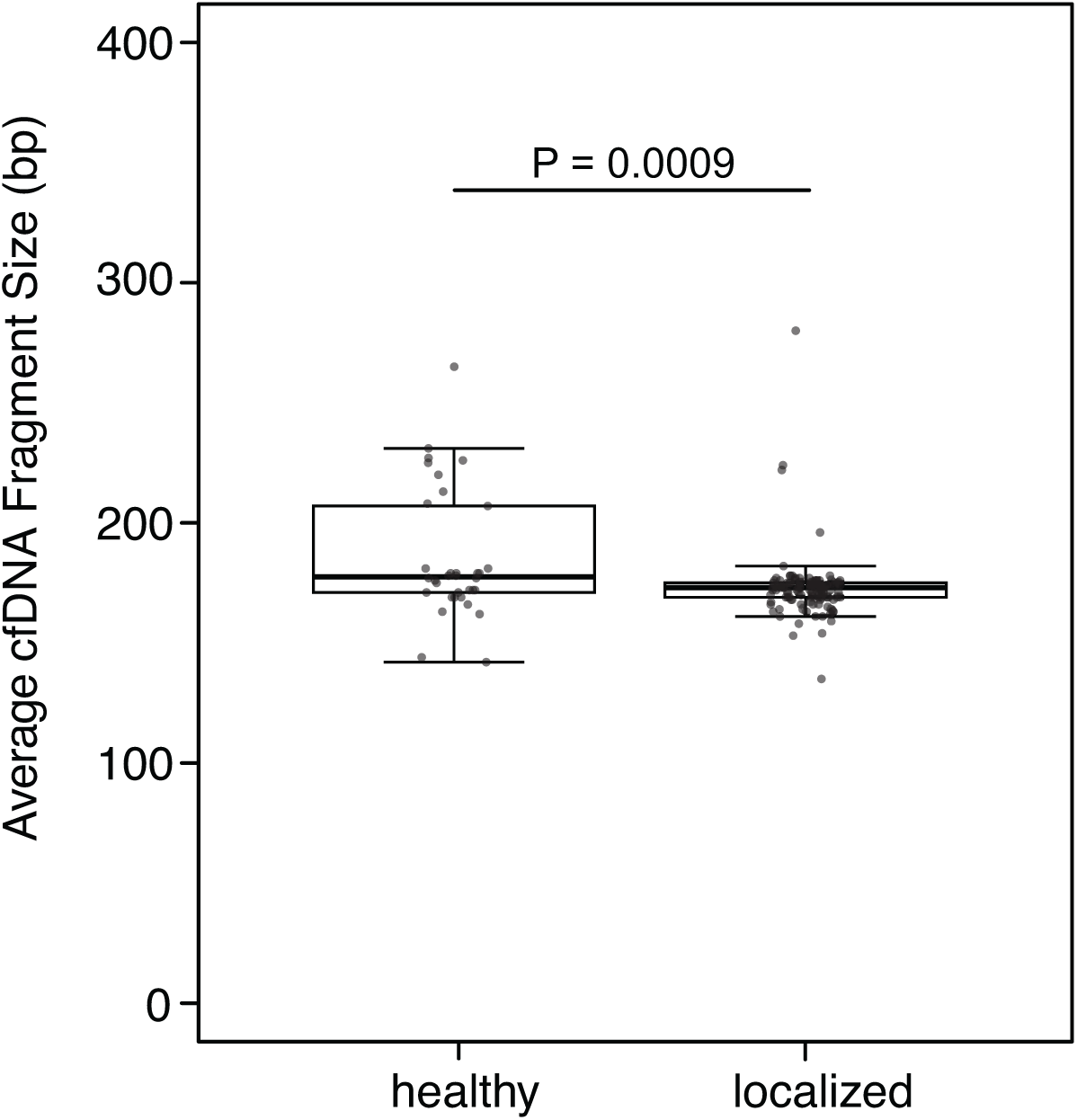
Distribution of average fragment size in healthy individuals and patients with localized disease (P = 0.0009). Boxplots and points identify the minimum, interquartile range, median, and maximum values for each group. A Mann-Whitney U-test was performed to test the difference in cfDNA fragment size.

### cfDNA Concentration / Fragment Size and Clinical Characteristics in Localized Prostate Cancer

There were no statistically significant differences in cfDNA concentration or fragment size for the clinical characteristics / outcomes evaluated here (Supplementary Table 2; Supplementary Table 3). Specifically, cfDNA concentration or fragment size did not appear to substantively differ across: pathologic Gleason score, organ confinement, extraprostatic extension, seminal vesicle invasion, time to biochemical recurrence, average cfDNA fragment size, clinical T stage, pathologic lymph node status, age at diagnosis, PSA at diagnosis, Decipher score, time to salvage therapy, and CAPRA-S score (Supplementary Figure 2; Supplementary Table 2; Supplementary Table 3). Additionally, no clear correlation was observed between cfDNA concentration and age at time of blood draw for healthy individuals, patients with localized disease, or patients with mCRPC (Supplementary Figure 3).

## Discussion

This study found that plasma cfDNA concentrations and fragment size may have diagnostic and prognostic value for detecting and profiling prostate cancer. Specifically, plasma cfDNA concentrations may help identify patients with advanced disease, while cfDNA fragment size may be used to distinguish patients with early stage disease from healthy individuals.

In the multivariate model that adjusted for age and PSA, plasma cfDNA concentration remained an independent predictor of mCRPC, indicating that cfDNA concentration may capture different biological processes than PSA and provides additional information (Table 2). Average cfDNA fragment size was predictive of localized disease. Looking at follow-up fragment size measures available for five patients, we found that one patient had a shorter average fragment size two months after surgery, and also exhibited post-treatment elevated PSA (Table 2; Supplementary Figure 6). In combination, these findings suggest that quantification of cfDNA overall may be valuable in identifying prostate cancer patients.

While the biological mechanisms underlying decreased fragment size in cancer patients are not well-understood, differences in nucleosome positioning and DNA methylation may result in varied DNA degradation. Our finding that the overall average fragment size in localized patients was shorter and more fragmented than in healthy individuals is consistent with the findings of studies assessing fragment size in patients with hepatocellular carcinoma and advanced pancreatic cancer^18,25^. The proportion of cfDNA fragments shorter than 150bp is also increased for multiple cancer types when compared to healthy fragment sizes with shallow genome-wide sequencing^17^. Localized prostate cancers are characterized by initial accumulation of clonal point mutations and deletions, with subsequent branching copy number gains where amplified regions are relatively enriched for tumor DNA, possibly modifying intracellular DNA degradation processes and mechanisms of DNA release and contributing to the size differences that were observed^26^.

The relatively short follow-up time for a protracted disease like prostate cancer is a limitation in this study. Of the 112 patients who underwent surgery, 24 patients experienced biochemical recurrence with a median follow-up time of three years (range, 9–1704 days), and it was not feasible to identify patients with localized disease who may have progressed to metastatic disease. This study did not include patients with metastatic disease who were not resistant to hormone therapy (i.e. castration sensitive), limiting the generalizability of these results to the full spectrum of this disease.

We processed samples in a manner that maximized the quality and quantity of extracted cfDNA^27–30^. An initial low-speed centrifugation step followed by high-speed centrifugation was used to reduce the amount of cellular debris and genomic DNA in the sample. Importantly, there was no significant difference in cfDNA concentration for samples collected in K3EDTA and PAXgene Blood ccfDNA tubes in the localized cohort (Supplementary Figure 5). However, the slightly increased cfDNA concentrations observed in controls may be due to cell lysis during transit, since whole blood was collected from individuals at a donor center in Kent, Washington and shipped overnight to San Francisco, California, whereas the patient samples were collected and processed onsite. Additionally, the lower overall cfDNA concentrations found in the localized cohort may be due to the subduing effect of anesthetic agents on cell death, which were administered prior to blood sample collection before surgery^31^. While data comparison across studies is difficult due to differences in sample collection and processing, most studies demonstrate the diagnostic role of cfDNA^12^. To quantify cfDNA, the Qubit 3.0 and the Agilent 2100 Bioanalyzer were used as a straightforward approach, albeit potentially less accurate than sequencing. In a clinical setting, an affordable, rapid, and straightforward test is critical to minimizing disruption to standard workflows while providing additional information. However, cfDNA quantification is a complementary approach that could help identify patients who may benefit from cfDNA sequencing.

Bastian et al. observed significant associations between cfDNA concentration and clinical characteristics in a cohort of patients with localized disease that experienced biochemical recurrence, supporting the hypothesis that cfDNA quantification may have more utility in the management of more advanced disease^14^. The lack of associations observed between cfDNA concentration and clinical characteristics in our study may be due to differences in the study cohorts. While all patients in the Bastian et al. study experienced BCR, only 24 of 112 patients experienced BCR in our study^14^.

A previous study demonstrated that in pre-treatment speciments, shorter cfDNA fragment size and elevated cfDNA concentrations were associated with shorter progression-free survival and overall survival in patients with advanced pancreatic cancer^25^. Due to the relatively short follow-up time in this study, future longitudinal studies evaluating disease progression from localized to metastatic disease are necessary to elucidate the value of analyzing cfDNA concentration and fragment size in the context of prostate cancer.

While the exact mechanism of cfDNA release into circulation remains unknown, apoptosis, lysis, necrosis, and active secretion have been identified as potential routes^6,32^. The cfDNA found in healthy individuals originates from hematopoietic cells, and likely reflects the processes of regulated cell turnover in these cells^8^. In patients with cancer, cfDNA includes both DNA fragments from hematopoietic cells, as well as fragments from tumor cells. Future studies evaluating the mechanisms of release will help elucidate the underlying biology of this biomarker, especially in combination with diagnostic and prognostic information over longer periods of time.

## Conclusion

Collectively, our data demonstrate the potential applications of plasma cfDNA concentration and cfDNA fragment size in both advanced and localized prostate cancer. Patients with advanced mCRPC had higher cfDNA concentration than men with localized disease or healthy controls, and those with localized disease had shorter average fragment sizes than controls. Importantly, cfDNA concentration and fragment size remained independent predictors after adjusting for age and PSA. Future studies assessing both cfDNA concentration and fragment size will be necessary to clarify the utility of plasma cfDNA in the context of diagnosis, prognosis, and disease monitoring.

## Acknowledgments

The authors would like to thank all participating patients and their families. This work was supported by the UCSF Goldberg-Benioff Program in Cancer Translational Biology, NIH R01 CA088164, NIH R01 CA201358, and the UCSF Moritz-Heyman Discovery Fellows Program.

## Supplementary Figures

**Supplementary Figure 1.**
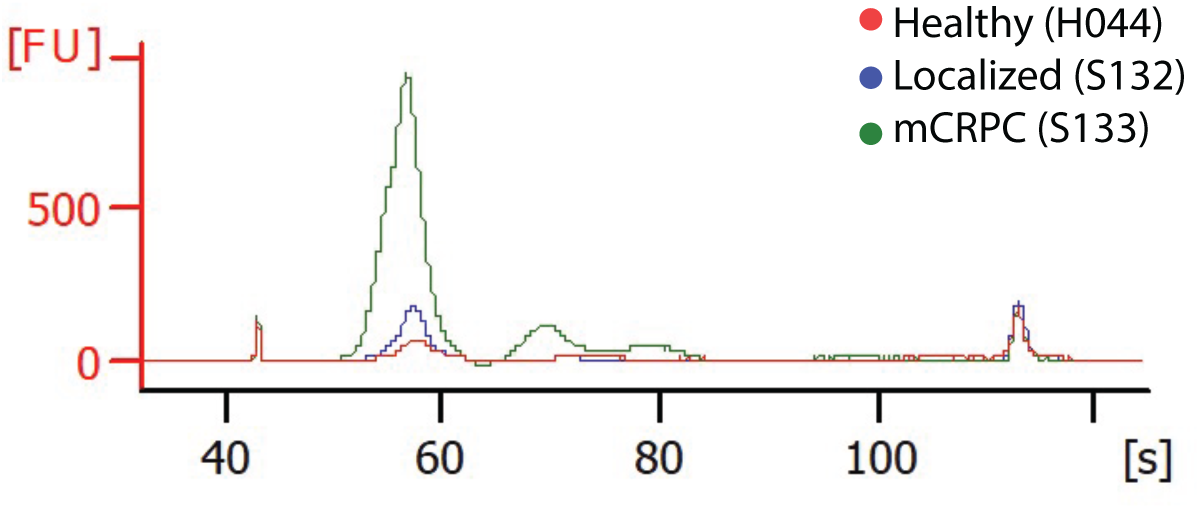
Plasma cfDNA concentration and distribution of cfDNA fragment size with representative traces for a healthy individual, patient with localized prostate cancer, and mCRPC patient.

**Supplementary Figure 2.**
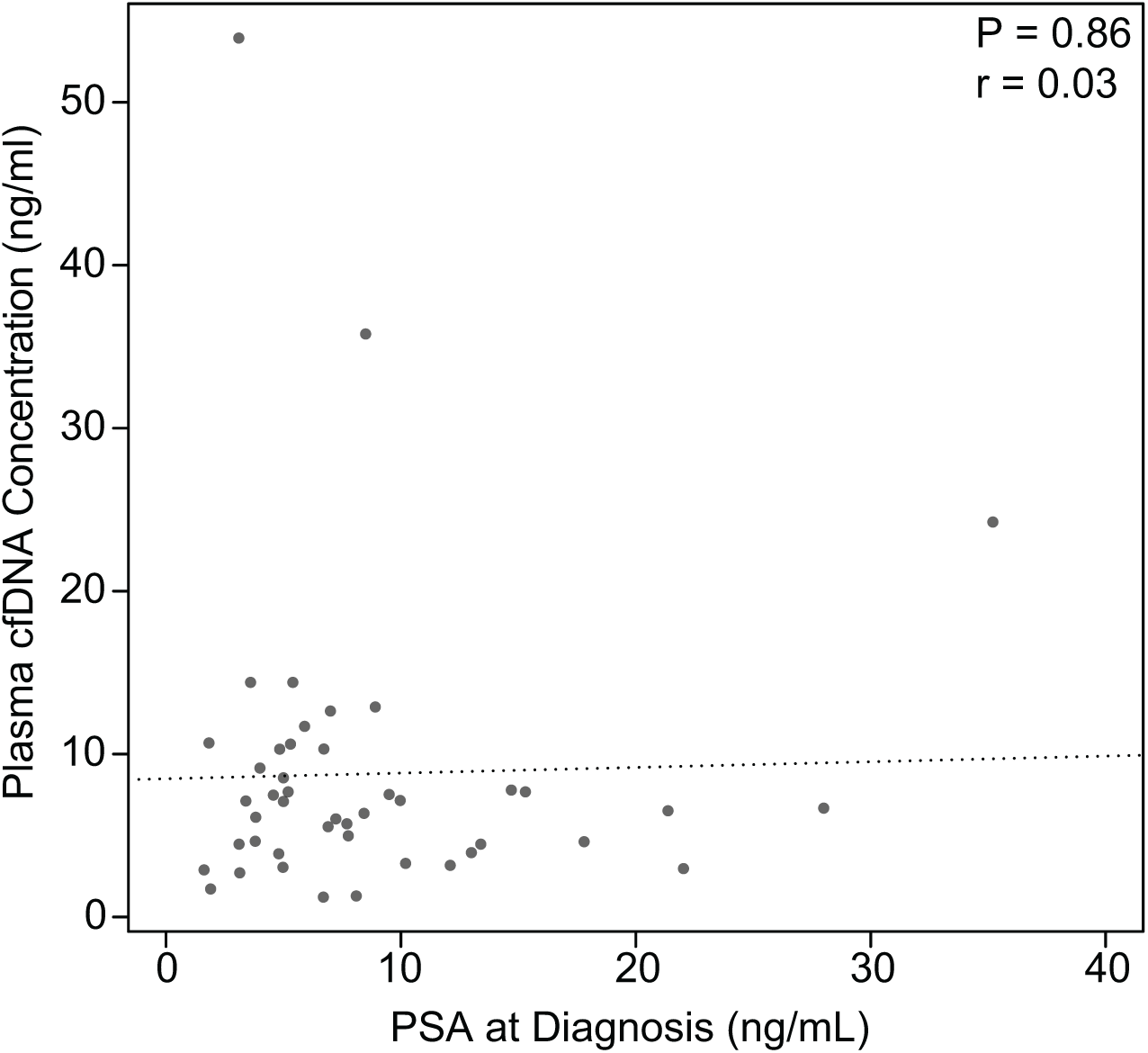
Assessment of the relationship between cfDNA concentration and PSA for patients with localized disease (P = 0.86).

**Supplementary Figure 3.**
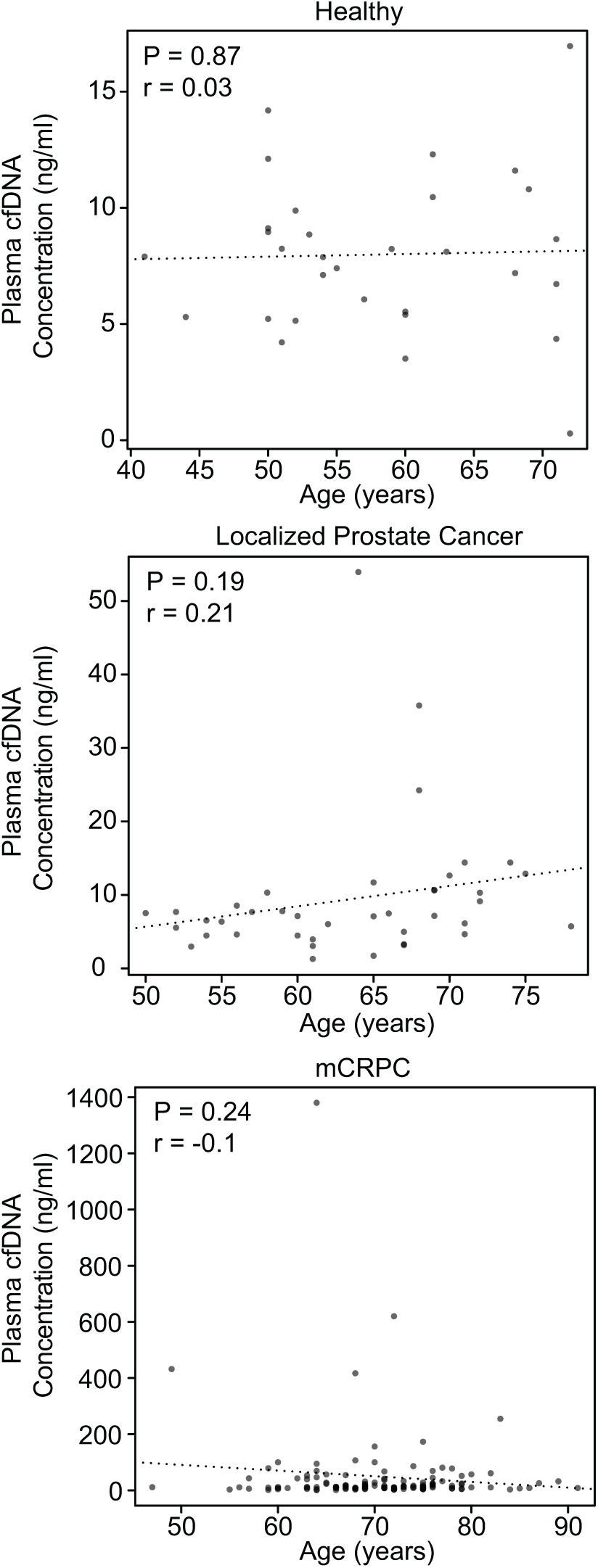
Assessment of the relationship between cfDNA concentration and age at time of blood draw.

**Supplementary Figure 4.**
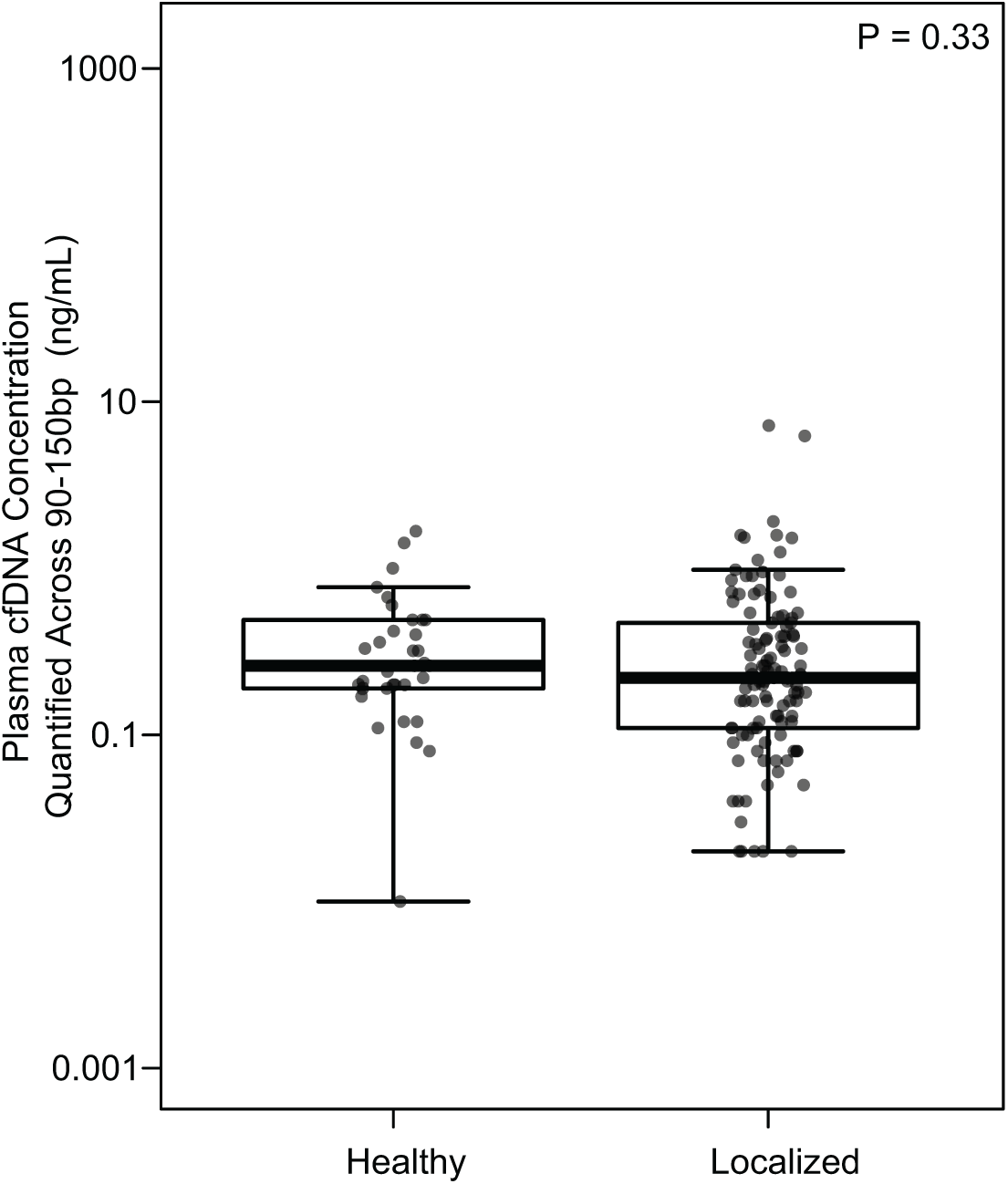
Distribution of plasma cfDNA concentration quantified across 90–150bp in healthy individuals and patients with localized disease with the 2100 Bioanalyzer. Boxplots and points identify the minimum, interquartile range, median, and maximum values for each group. A Mann-Whitney U-test was performed to test the difference in cfDNA yields.

**Supplementary Figure 5.**
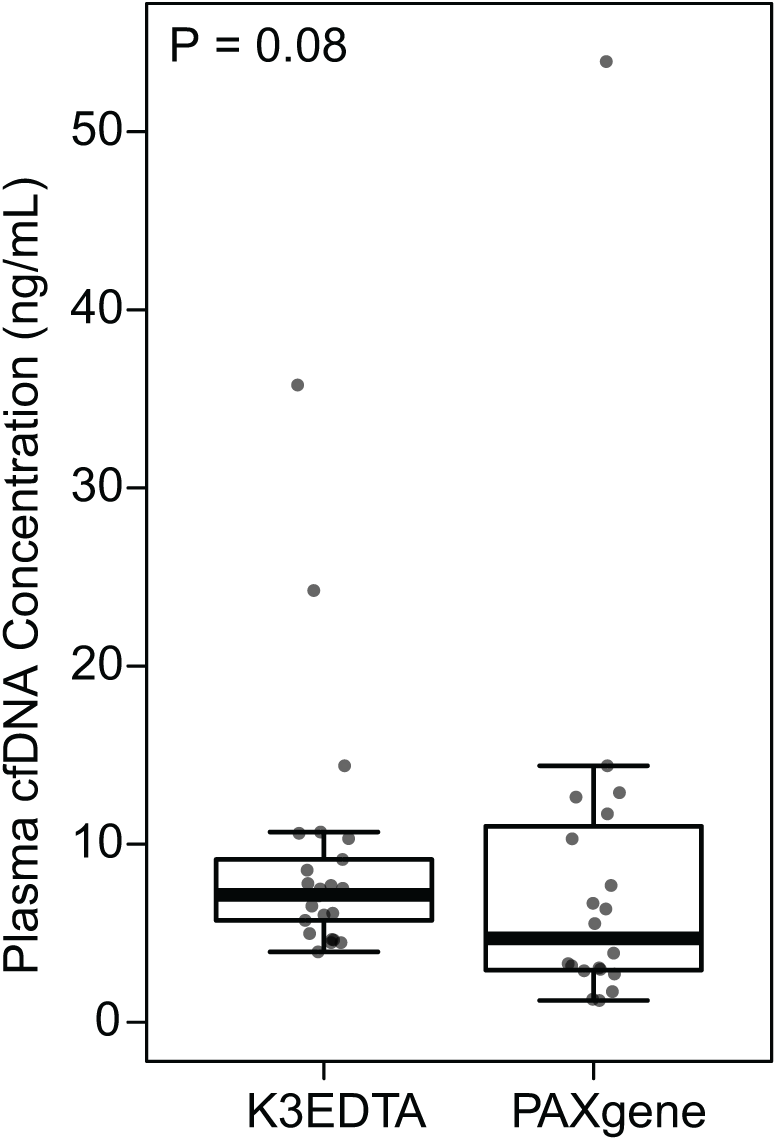
Distribution of cfDNA concentration for K3EDTA and Qiagen PAXgene tube types for patients with localized disease.

**Supplementary Figure 6.**
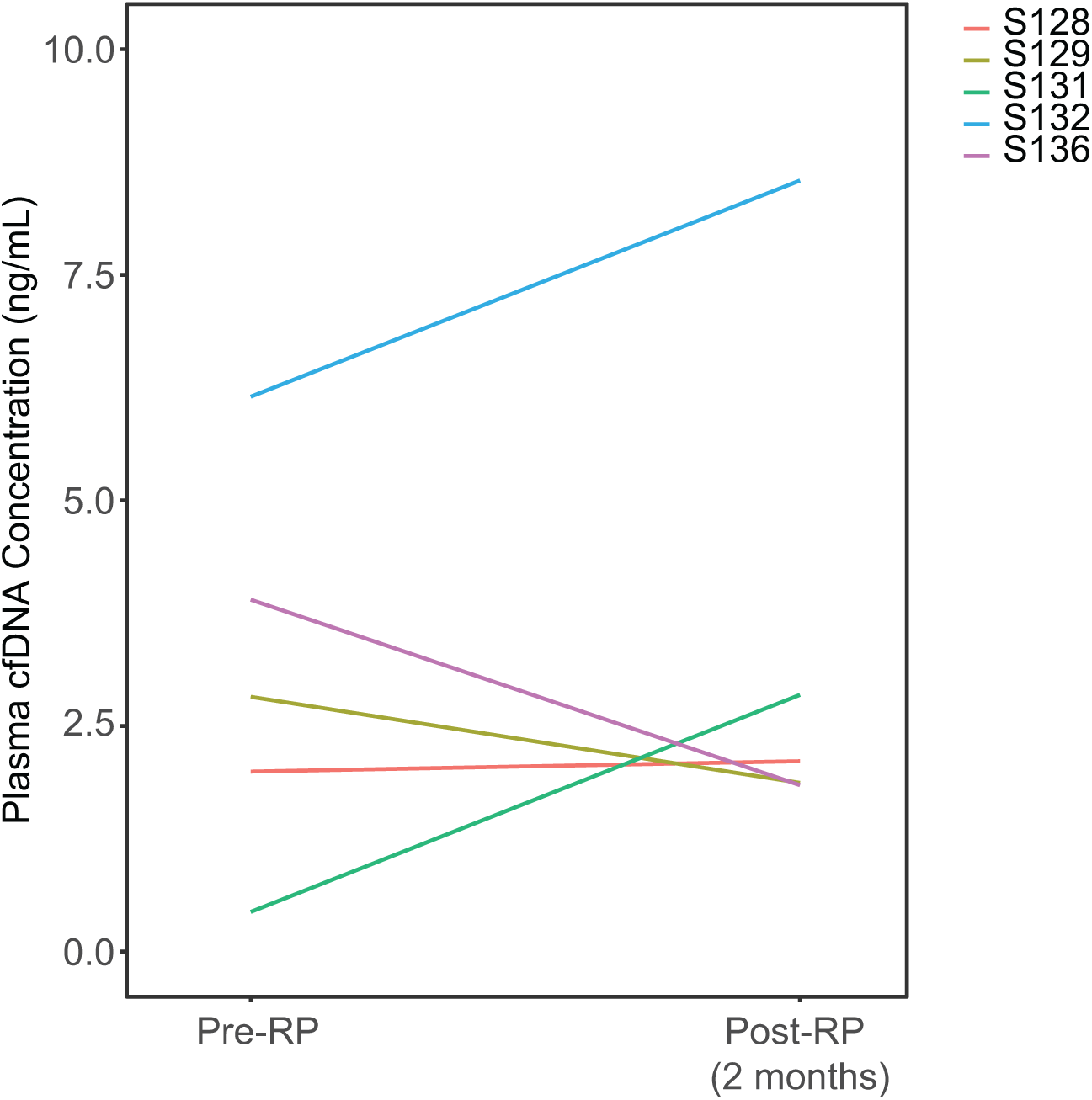
Blood samples were collected two months after RP for five patients. Plasma cfDNA concentration increased for two patients (S131 and S132). Patient S131 exhibited a decrease in average cfDNA fragment size from 206bp to 167bp, as well as an elevated PSA of 0.36 ng/mL 64 days after surgery, while the average cfDNA fragment size for patient S132 remained the same at 181bp and PSA levels remained low at 0.03 ng/mL.

**Supplementary Table 1.**
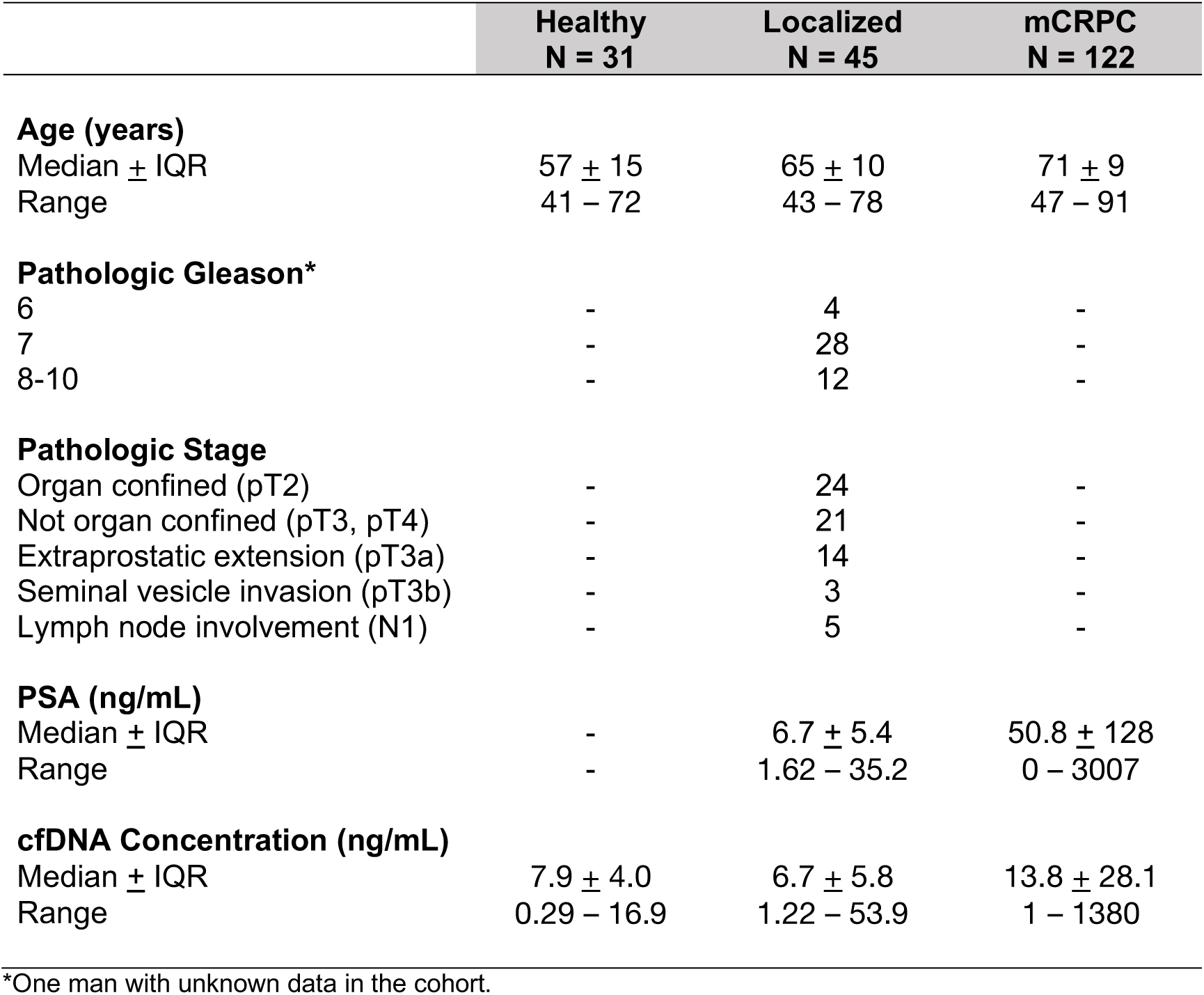
Clinical characteristics of individuals included in the cfDNA concentration analysis.

**Supplementary Table 2.** Association between log transformed cfDNA concentration or average cfDNA fragment size and categorical clinical features for patients with localized disease.

| Clinical Feature | Number of Patients | Average cfDNA Concentration (ng/mL) | Average cfDNA Fragment Size (bp) |
| --- | --- | --- | --- |
| Pathologic Gleason score |  |  |  |
| $\leq 7$ (3+4) | 25 | 8.90 | 169.8 |
| $\geq 7$ (4+3) | 20 | 8.65 | 174.7 |
| P |  | 0.19 | 0.09 |
| Organ confinement |  |  |  |
| Confined (pT2) | 24 | 8.25 | 172.7 |
| Not confined (pT3, pT4) | 21 | 9.40 | 172.8 |
| P |  | 0.80 | 0.69 |
| Extraprostatic extension |  |  |  |
| Negative (pT2) | 24 | 8.25 | 172.7 |
| Positive(pT3a) | 14 | 12.33 | 170.7 |
| P |  | 0.29 | 0.67 |
| Seminal vesicle invasion |  |  |  |
| Negative (pT2, pT3a) | 38 | 9.75 | 171.7 |
| Positive (pT3b) | 3 | 3.74 | 171.9 |
| P |  | 0.15 | 0.45 |
| Pathologic lymph node |  |  |  |
| Negative (pN0) | 23 | 8.45 | 172.2 |
| Positive (pN1) | 5 | 8.49 | 173.8 |
| P |  | 0.90 | 0.69 |
| Biochemical Recurrence |  |  |  |
| No | 37 | 7.62 | 172.5 |
| Yes | 5 | 20.3 | 171.5 |
| P |  | 0.24 | 0.41 |
| Clinical T stage |  |  |  |
| T1c* | 20 | 9.47 | 169.5 |
| T2a-c<sup>†</sup> | 21 | 9.19 | 174.1 |
| P |  | 0.24 | 0.29 |

\* Tumor identified by needle biopsy, but clinically inapparent and not palpable
<sup>†</sup> Tumor is palpable

**Supplementary Table 3.** Association between cfDNA and continuous clinical features for patients with localized disease.

| Clinical Feature |  | cfDNA<br>Concentration | Average<br>Fragment Size |
| --- | --- | --- | --- |
| Age at Diagnosis (years) |  |  |  |
| Number of Patients | 45 |  |  |
| Average (SD) | 62.7 (7) |  |  |
| P |  | 0.35 | 0.60 |
| PSA at Diagnosis (ng/mL) |  |  |  |
| Number of Patients | 45 |  |  |
| Average (SD) | 8.7 (7) |  |  |
| P |  | 0.86 | 0.85 |
| Decipher Score |  |  |  |
| Number of Patients | 12 |  |  |
| Average (SD) | 0.66 (0.2) |  |  |
| P |  | 0.81 | 0.65 |
| Time to Salvage Therapy (days) |  |  |  |
| Number of Patients | 39 |  |  |
| Average (SD) | 1044 (926) |  |  |
| P |  | 0.63 | 0.25 |
| Average Fragment Size (bp) |  |  |  |
| Number of Patients | 45 |  |  |
| Average (SD) | 176 (21) |  |  |
| P |  | 0.67 | - |
| CAPRA-S Score |  |  |  |
| Number of Patients | 45 |  |  |
| Average (SD) | 3.7 (3) |  |  |
| P |  | 0.50 | 0.12 |

## References

1. Siegel, R. L., Miller, K. D. & Jemal, A. Cancer statistics, 2019. CA. Cancer J. Clin. 69, 7–34 (2019).

2. Lilja, H., Ulmert, D. & Vickers, A. J. Prostate-specific antigen and prostate cancer: Prediction, detection and monitoring. Nat. Rev. Cancer 8, 268–278 (2008).

3. Carroll, P. H. & Mohler, J. L. NCCN guidelines updates: Prostate cancer and prostate cancer early detection. JNCCN J. Natl. Compr. Cancer Netw. 16, 620–623 (2018).

4. Mandel, P. & Metais, P. Les acides nucleiques du plasma sanguin ches l’homme. C R Seances Soc Biol Fil. 142, 241–243 (1948).

5. Stroun, M. et al. Neoplastic characteristics of the DNA found in the plasma of cancer patients. Oncology 46, 318–22 (1989).

6. Stroun, M., Lyautey, J., Lederrey, C., Olson-Sand, A. & Anker, P. About the possible origin and mechanism of circulating DNA: Apoptosis and active DNA release. Clin. Chim. Acta 313, 139–142 (2001).

7. Jahr, S. et al. DNA fragments in the blood plasma of cancer patients: Quantitations and evidence for their origin from apoptotic and necrotic cells. Cancer Res. 61, 1659–1665 (2001).

8. Lui, Y. Y. N. et al. Predominant hematopoietic origin of cell-free dna in plasma and serum after sex-mismatched bone marrow transplantation. Clin. Chem. 48, 421–427 (2002).

9. Koffler, D., Agnello, V., Winchester, R. & Kunkel, H. G. The occurrence of single-stranded DNA in the serum of patients with systemic lupus erythematosus and other diseases. J. Clin. Invest. 52, 198–204 (1973).

10. Tan, E. M., Schur, P. H., Carr, R. I. & Kunkel, H. G. Deoxybonucleic acid (DNA) and antibodies to DNA in the serum of patients with systemic lupus erythematosus. J. Clin. Invest. 45, 1732–1740 (1966).

11. Tissot, C. et al. Circulating free DNA concentration is an independent prognostic biomarker in lung cancer. Eur Respir J 46, 1773–1780 (2015).

12. Ellinger, J. et al. The role of cell-free circulating DNA in the diagnosis and prognosis of prostate cancer. Urol. Oncol. Semin. Orig. Investig. 29, 124–129 (2011).

13. Jung, K. et al. Increased cell-free DNA in plasma of patients with metastatic spread in prostate cancer. Cancer Lett. 205, 173–80 (2004).

14. Bastian, P. J. et al. Prognostic value of preoperative serum cell-free circulating DNA in men with prostate cancer undergoing radical prostatectomy. Clin. Cancer Res. 13, 5361–5367 (2007).

15. Feng, J. et al. Plasma cell-free DNA and its DNA integrity as biomarker to distinguish prostate cancer from benign prostatic hyperplasia in patients with increased serum prostate-specific antigen. Int Urol Nephrol 45, 1023–1028 (2013).

16. Lapin, M. et al. Fragment size and level of cell-free DNA provide prognostic information in patients with advanced pancreatic cancer. J. Transl. Med. 16, 1–10 (2018).

17. Mouliere, F. et al. Enhanced detection of circulating tumor DNA by fragment size analysis. Sci. Transl. Med 10, (2018).

18. Jiang, P. et al. Lengthening and shortening of plasma DNA in hepatocellular carcinoma patients. Proc. Natl. Acad. Sci. U. S. A. 112, E1317–E1325 (2015).

19. Underhill, H. R. et al. Fragment Length of Circulating Tumor DNA. 1–24 (2016). doi: 10.1371/journal.pgen.1006162

20. Marrone, M., Potosky, A. L., Penson, D. & Freedman, A. N. A 22 gene-expression assay, decipher® (GenomeDx biosciences) to predict five-year risk of metastatic prostate cancer in men treated with radical prostatectomy. PLoS Curr. 7, 1–8 (2015).

21. Cooperberg, M. R., Hilton, J. F. & Carroll, P. R. The CAPRA-S score: a straightforward tool for improved prediction of outcomes after radical prostatectomy. Cancer 117, 5039–5046 (2011).

22. Cooperberg, M. R. et al. Combined value of validated clinical and genomic risk stratification tools for predicting prostate cancer mortality in a high-risk prostatectomy cohort. Eur. Urol. 67, 326–333 (2015).

23. Liu, H. et al. Prognostic significance of six clinicopathological features for biochemical recurrence after radical prostatectomy: A systematic review and meta-analysis. Oncotarget 9, 32238–32249 (2018).

24. Wyatt, A. W. et al. Concordance of Circulating Tumor DNA and Matched Metastatic Tissue Biopsy in Prostate Cancer. doi: 10.1093/jnci/djx118

25. Lapin, M. et al. Fragment size and level of cell-free DNA provide prognostic information in patients with advanced pancreatic cancer. J. Transl. Med. 16, 300 (2018).

26. Melijah, S. et al. The Evolutionary Landscape of Localized Prostate Cancers Drives Clinical Aggression. Cell 173, (2018).

27. Meddeb, R., Pisareva, E. & Thierry, A. R. Guidelines for the preanalytical conditions for analyzing circulating cell-free DNA. Clin. Chem. 65, 623–633 (2019).

28. Lee, T.-H., Montalvo, L., Chrebtow, V. & Busch, M. P. Quantitation of genomic DNA in plasma and serum samples: higher concentrations of genomic DNA found in serum than in plasma. Transfusion 41, 276–282 (2001).

29. Jung, M., Klotzek, S., Lewandowski, M., Fleischhacker, M. & Jung, K. Changes in concentration of DNA in serum and plasma during storage of blood samples [5]. Clin. Chem. 49, 1028–1029 (2003).

30. Devonshire, A. S. et al. Towards standardisation of cell-free DNA measurement in plasma: Controls for extraction efficiency, fragment size bias and quantification. Anal. Bioanal. Chem. 406, 6499–6512 (2014).

31. Prakash, K., Aggarwal, S., Bhardwaj, S., Ramakrishna, G. & Pandey, C. K. Serial perioperative cell-free DNA levels in donors and recipients undergoing living donor liver transplantation. Acta Anaesthesiol. Scand. 61, 1084–1094 (2017).

32. Sozzi, G. et al. Quantification of free circulating DNA as a diagnostic marker in lung cancer. J. Clin. Oncol. 21, 3902–3908 (2003).

